# CLEAR-HPV: Interpretable Concept Discovery for HPV-Associated Morphology in Whole-Slide Histology

**DOI:** 10.64898/2026.02.04.703870

**Authors:** Weiyi Qin, Yingci Liu-Swetz, Shiwei Tan, Hao Wang

**Affiliations:** Department of Computer Science, Rutgers University, New Brunswick, NJ, USA; Rutgers Health, Rutgers University, Newark, NJ, USA

**Keywords:** computational pathology, multiple instance learning, interpretable deep learning, concept discovery, head and neck cancer, human papillomavirus (HPV), whole slide image analysis

## Abstract

Human papillomavirus (HPV) status is a critical determinant of prognosis and treatment response in head and neck and cervical cancers. Although attention-based multiple instance learning (MIL) achieves strong slide-level prediction for HPV-related whole-slide histopathology, it provides limited morphologic interpretability. To address this limitation, we introduce Concept-Level Explainable Attention-guided Representation for HPV (CLEAR-HPV), a framework that restructures the MIL latent space using attention to enable concept discovery without requiring concept labels during training. Operating in an attention-weighted latent space, CLEAR-HPV automatically discovers keratinizing, basaloid, and stromal morphologic concepts, generates spatial concept maps, and represents each slide using a compact concept-fraction vector. CLEAR-HPV’s concept-fraction vectors preserve the predictive information of the original MIL embeddings while reducing the high-dimensional feature space (e.g., 1536 dimensions) to only 10 interpretable concepts. CLEAR-HPV demonstrates consistent concept structure across TCGA-HNSCC, TCGA-CESC, and CPTAC-HNSCC, providing compact, concept-level interpretability through a general, backbone-agnostic framework for attention-based MIL models of whole-slide histopathology. All original code, preprocessing scripts, and trained model checkpoints are available on GitHub (https://github.com/Wang-ML-Lab/CLEAR-HPV))

## INTRODUCTION

HPV-associated head and neck and cervical cancers together account for 690,000 new cases worldwide each year^1^, with HPV status strongly stratifying survival, treatment intensity, and long-term functional outcomes, making accurate and interpretable HPV assessment a problem of major clinical and public health significance. HPV-positive tumors are often non-keratinizing or basaloid^2–4^, whereas HPV-negative tumors more commonly display keratinizing squamous morphology^2,3,5^. However, substantial morphologic overlap and variability mean that HPV status cannot be reliably inferred by human observers from routine histologic assessment alone^6^; ancillary immunohistochemical or molecular assays therefore remain the standard of care^7,8^, but they often incur substantially higher costs, owing to additional reagents, instrumentation, and technical processing.

On the other hand, digital histology (and histopathology) and the use of whole slide images (WSIs) have recently shown promising accuracy in capturing nuanced patterns that human observers often miss; however, they are often lacking in interpretability, limiting their clinical adoption. As a result, there is a pressing need for the best of both worlds: computational histology methods that not only classify slides accurately but also reveal how predictions relate to recognizable, biologically grounded morphologic patterns, with stability across different staining, scanning, and institutional settings.

Recent deep learning methods, including vision foundation models, have achieved strong performance in WSI classification across diverse diagnostic and molecular tasks^7,9–11^, but most models function as black boxes that offer limited interpretability on what histologic patterns are driving their predictions^12–14^. Weakly supervised multiple instance learning (MIL) is a widely adopted framework for WSI analysis, and popular MIL models (e.g., ABMIL^15^, CLAM^16^, Trans-MIL^17^) treat each slide as a large, heterogeneous collection of tiles with only slide-level labels. Widely used interpretability methods such as attention heatmaps and Grad-CAM^18^ indicate where a model attends, but do not provide concept-level, human-understandable explanations of what histologic patterns are driving its predictions. As a result, current approaches offer only coarse, qualitative cues and cannot identify the discrete, reproducible morphologic concepts present in WSIs. This limits biological insight, reduces reproducibility across sites, and weakens trust in model predictions, therefore limiting clinical deployment.

These interpretability limitations motivate a closer examination of how MIL models internally encode morphology to determine whether their representations can be reorganized into *clinically meaningful and human-understandable concepts*. Prior work has shown that deep neural networks naturally organize intermediate features into latent spaces that encode semantic or visual factors^19,20^. In attention-based MIL models, the tile-level embeddings produced before attention pooling define an intermediate *h*-space that captures the morphologic features learned by the model across tumor and stromal regions. Latent spaces such as the *h*-space can be reorganized into human-interpretable concepts^21^, and concept discovery from neural embeddings has been shown to recover coherent visual structures^22–26^. Together, these observations suggest that MIL backbones already encode rich morphologic structure, but require an attention-aware organization strategy to make this structure explicit and biologically interpretable.

In this work, we show that attention mechanisms in MIL induce a latent morphologic structure that can be reorganized into discrete histologic concepts without tile-level annotations. We develop CLEAR-HPV (Concept-Level Explainable Attention-guided Representation for HPV), a framework that restructures the attention-weighted MIL *h*-space to enable *annotation-free concept discovery* in HPV-related histopathology. Rather than modifying the classifier or explicitly optimizing for higher accuracy, CLEAR-HPV operates *post hoc* on the latent embeddings of a *trained* model (e.g., CLAM^16^), using attention weights to focus concept discovery on tiles the model already considers informative. The framework yields coherent keratinizing, basaloid, and stromal morphologic concepts that align with established HPV-associated patterns, together with two complementary interpretable outputs, *spatial concept maps* that show where concepts appear across each slide, and *compact concept-fraction representations* that summarize slide-level tissue composition in a quantitative, low-dimensional form.

Applied to three cohorts of data, TCGA-HNSCC^27^, TCGA-CESC^27^, and CPTAC-HNSCC^28^, CLEAR-HPV discovers stable concepts and consistent concept-fraction patterns that generalize across cohorts, indicating that the discovered morphology reflects consistent HPV-associated structure rather than dataset-specific artifacts. The resulting concept-fraction representations preserve the discriminative structure encoded in the original MIL embeddings, allowing down-stream classifiers to recover comparable slide-level predictions while operating on an interpretable concept space — achieving the best of both worlds. Together, these findings demonstrate that the latent feature space of attention-based MIL already contains rich, biologically meaningful organization, and that our attention-guided concept discovery can expose this structure without sacrificing predictive performance. Figure 1 shows an overview of our CLEAR-HPV; implementation details are described in the Methods section.

**Figure 1:**
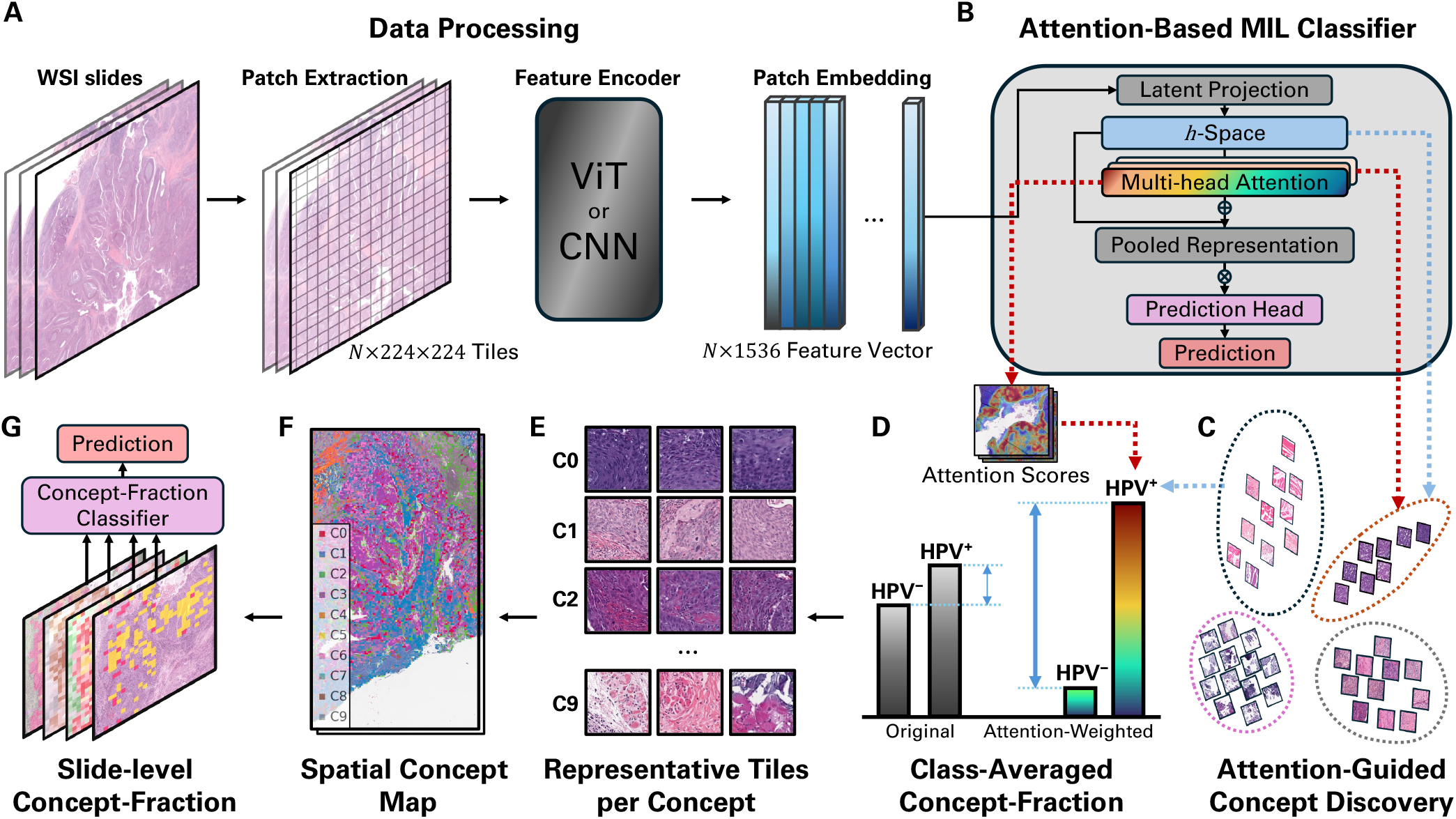
Overview of the CLEAR-HPV framework. **(A)** Data processing pipeline: WSIs are decomposed into fixed-size tiles, encoded with a pretrained ViT or CNN, and converted into patch-level feature embeddings. **(B)** An attention-based MIL classifier projects embeddings into the *h*-space latent representation and uses multi-head attention to compute tile-level contributions, which are pooled into a single slide-level embedding for HPV prediction. **(C)** CLEAR-HPV performs annotation-free concept discovery on attention-weighted *h*-space embeddings to identify coherent morphologic concepts. **(D)** Using the discovered concepts, each slide is represented by a concept-fraction vector, which is then averaged across slides to obtain class-averaged concept-fraction vectors that summarize morphologic composition for HPV-positive and HPV-negative cohorts. **(E)** Representative tiles illustrate the characteristic morphology captured by each discovered concept. **(F)** Spatial concept maps visualize the distribution of concepts across the WSIs, revealing their spatial organization. **(G)** Slide-level concept-fraction vectors provide an interpretable representation that supports a concept-fraction classifier, which recovers MIL predictive performance while offering concept-level explanations. More details are available in the Methods section.

In summary, our primary contributions are as follows:

1. We introduce CLEAR-HPV, the first general framework to automatically discover pathology-relevant morphologic concepts for HPV prediction without tile-level supervision (or annotation).
2. We demonstrate that CLEAR-HPV leverages the attention-weighted latent space in a deep learning model to produce spatial concept maps and concept-fraction vectors, offering biologically grounded, concept-level interpretability for whole-slide histopathology.
3. CLEAR-HPV preserves the predictive performance of the interpreted model while reducing its high-dimensional features (e.g., 1536 dimensions) to only 10 interpretable concepts.
4. We show that CLEAR-HPV is compatible with diverse attention-based MIL backbones and retains strong slide-level performance across diverse architectural designs, enabling robust and consistent concept discovery beyond a single model instantiation.
5. We further show that these concept-level representations are stable across different cohorts (e.g., TCGA and CPTAC), preserve clinically relevant predictive signal under transfer, and reveal cross-cohort consistency of HPV-related morphology.

## RESULTS

In this study, we evaluated CLEAR-HPV across three independent WSI cohorts (datasets) to examine whether biologically coherent and interpretable concepts can be discovered across diverse clinical and technical settings. The TCGA-HNSCC cohort included 102 patients and 106 diagnostic WSIs (38 HPV-positive, 64 HPV-negative). The TCGA-CESC cohort consisted of 146 patients and 154 WSIs, predominantly HPV-positive (138 HPV-positive, 8 HPV-negative)^27^. The CPTAC-HNSCC dataset contributed 112 HPV-negative patients and 368 WSIs, serving as an external validation cohort collected under a different study protocol^28^. These datasets differ substantially in HPV prevalence, staining and scanning protocols, and clinical outcomes, allowing us to assess whether CLEAR-HPV identifies stable morphologic concepts rather than cohort-specific artifacts.

CLEAR-HPV is designed as a post-hoc explainability framework; its goal is *not* to improve predictive accuracy but to discover and interpret the morphologic concepts encoded in the model’s internal representations. Compared to the interpreted (explained) deep learning model that uses *high-dimensional* (e.g., 1536 dimensions), *uninterpretable* embeddings, CLEAR-HPV discovers a *compact, interpretable* set of concepts, e.g., only 10 concepts. Therefore, results are considered *very strong* as long as *comparable* predictive performance (e.g., AUC, ACC, F1) can be achieved using CLEAR-HPV’s discovered concepts.

We use CLAM^16^, a widely adopted attention-based multiple instance learning (MIL) method, as the primary target backbone (base) model to explain. CLAM provides tile-level attention scores and a learned intermediate latent space (the *h*-space). Together, these form the foundation for concept discovery. We also provide results on three other backbone models and their variants to demonstrate the generality of CLEAR-HPV (Table 3).

A consistent 10-fold protocol was applied to each cohort, enabling systematic assessment of interpretability and robustness across heterogeneous datasets (Figure 1).

### Base model performance

Table 1 shows that, on TCGA-HNSCC, the CLAM attention-based MIL backbone achieved consistent slide-level performance (ACC = 0.77 *±* 0.06, AUC = 0.86 *±* 0.05), indicating that its learned representations capture generalizable, discriminative structure.

**Table 1:** Cross-cohort generalization of the baseline CLAM model. ^a^ AUC, Precision, Recall, and F1 are not applicable (N/A) because the CPTAC-HNSCC cohort contains only HPV-negative slides (single-class).

| Dataset | ACC | AUC | Prec | Rec | F1 |
| --- | --- | --- | --- | --- | --- |
| TCGA-HNSCC | 0.765±0.063 | 0.863±0.051 | 0.788±0.098 | 0.673±0.145 | 0.696±0.101 |
| TCGA-CESC | 0.593±0.083 | 0.684±0.154 | 0.977±0.029 | 0.581±0.176 | 0.716±0.073 |
| CPTAC-HNSCC<sup>a</sup> | 0.704±0.120 | N/A | N/A | N/A | N/A |

We then evaluated the model in a fully zero-shot setting on external cohorts *without any calibration or retraining*. On TCGA-CESC, a cohort dominated by HPV-positive tumors, performance decreased as expected due to differences in tissue origin and histomorphologic context (AUC ≈ 0.68), but the model retained very high precision (≈0.98), indicating preservation of class-specific structure under domain shift. On CPTAC-HNSCC, where only HPV-negative cases are available for evaluation, accuracy remained consistent (0.70 *±* 0.12), demonstrating robustness to moderate staining and scanner variability. Together, these zero-shot evaluations show that the CLAM backbone maintains consistent decision behavior across cohorts and preserves transferable class-related structure in its latent representation, making it suitable for downstream concept-level analysis.

### Annotation-free concept discovery in the *h*-space

CLEAR-HPV is motivated by the observation that the latent *h*-space of attention-based MIL models contains rich morphologic structure that can be made explicit via concept discovery. Here, the *h*-space consists of tile-level embeddings produced before attention pooling, illustrated as a blue box in Figure 1(B). The attention weights learned by the backbone identify which regions contribute most to the final prediction, and attention pooling is typically used to pool these tile-level embeddings into a single embedding vector (i.e., “Pooled Representation” in Figure 1(B)), which is fed into a prediction head for classification. CLEAR-HPV constructs an attention-guided representation by weighting each embedding *h*_*i*_ by its attention score *α*_*i*_, thereby emphasizing diagnostically informative tiles while reducing background variation. More details are provided in the Methods section.

#### Concept-discovery methods and baselines

To evaluate the discovered concepts, all concept discovery methods were applied to the TCGA-HNSCC training folds, which provide sufficient morphologic diversity for assessing cluster structure. Concept discovery was performed with *K* = 10 concepts. This choice is supported by consistent empirical evidence: Table S1 shows that predictive performance remains stable across *K* ∈ {5, 10, 15} with overlapping confidence intervals across all major metrics, indicating that *K* = 10 achieves comparable performance without sensitivity to the exact choice of *K*. Table S2 further demonstrates that concept geometry is highly stable across resolutions, with forward persistence and reverse fragmentation both exceeding 0.96, confirming that the discovered concepts are preserved under changes in *K*. The elbow analysis (Figure S1) provides additional support, showing diminishing returns in clustering compactness beyond *K* = 10. We evaluate two variants of CLEAR-HPV: concepts produced from raw-*h*-space embeddings, i.e., “CLEAR-HPV (raw-*h*)”, and concepts derived from attention-weighted (AW) *h*-space embeddings, i.e., “CLEAR-HPV (AW-*h*)”. These form the primary concept sets used throughout the analysis. For comparison, we evaluated several baseline methods that differ in where and how morphologic structure is extracted. Specifically, we considered (more details in the Methods section): (1) heatmap-based grouping, which reflects which tiles the model attends to, without defining discrete concept distributions, (2) encoder-feature clustering, which generates unconstrained concept distributions derived from encoder feature representations, and (3) a Dirichlet concept model, which represents concept membership probabilistically and allows tiles and slides to express varying degrees of concept mixing. Together, these baselines compare different input representations and concept construction approaches, highlighting how attention-structured *h*-space representations influence the coherence and interpretability of discovered morphologic concepts.

#### Quantitative comparison of concept-discovery methods

After defining the set of concept-discovery methods, we evaluate all approaches under a common framework to assess how well the resulting concepts summarize slide-level morphology and preserve diagnostically relevant signals. With *K* concepts in total, each slide is summarized by a *K*-dimensional *concept-fraction vector* that quantifies the proportion of tiles assigned to each concept. The concept-fraction vector is the core representation used throughout our analysis, providing a unified and interpretable summary of slide-level morphology derived from tile-level concepts. To assess whether these concepts capture diagnostically meaningful information, we introduce a *concept-fraction classifier* that maps concept fraction vectors to HPV status, without introducing additional trainable parameters.

A detailed description is provided in the Methods section. We use common classification metrics such as Accuracy (ACC), Area Under the Curve (AUC), Precision, Recall, and F1 measure how well each representation preserves the predictive signal present in the original MIL embeddings. Note that our goal is *not* to improve accuracy upon the original MIL backbone (e.g., CLAM). Instead, we aim to measure how well the concept-based explanations can retain the predictive performance of the original MIL backbone for HPV status. Therefore we consider these results strong when the CLEAR-HPV predictor achieves performance (e.g., ACC or AUC) comparable to the original MIL backbone.

Table 2 shows the performance of classifying HPV for all concept-discovery methods using only the discovered concept-fraction vectors, evaluated with a simple rule-based concept-fraction classifier. Our CLEAR-HPV’s two variants, raw-*h* and AW-*h*, capture complementary structure. Raw-*h* preserves the intrinsic latent space learned by the MIL encoder and achieves the highest AUC, while AW-*h* produces more coherent and stable morphologic concepts by amplifying high-attention tiles. Notably, even using only the discovered concept-fraction vector (with only *K* = 10 dimensions), without access to the original tile embeddings (with 1536 dimensions), both variants retain predictive performance comparable to the CLAM backbone, indicating that the concept-fraction representation preserves the discriminative signal of the original model despite substantial dimensionality reduction.

**Table 2:** Comparison of concept-discovery methods on TCGA-HNSCC. The heatmap provides an attention-only reference and does not define discrete concepts. Encoder concepts derive clusters directly from encoder feature representations. Within the *h*-space, Dirichlet concepts serve as an alternative concept discovery baseline applied to the unweighted latent space. In contrast, CLEAR-HPV performs concept discovery directly in the attention-weighted latent *h*-space. Metrics assess whether the resulting concept-fractions vectors retain the predictive signal present in the original MIL backbone. Results on more metrics are provided in Table S3.

| Method | ACC | AUC | Prec | Rec | F1 |
| --- | --- | --- | --- | --- | --- |
| Heatmap | 0.538±0.110 | 0.674±0.147 | 0.503±0.137 | 0.606±0.215 | 0.538±0.114 |
| Dirichlet Concepts | 0.646±0.071 | 0.641±0.134 | 0.475±0.314 | 0.200±0.208 | 0.266±0.193 |
| Encoder Concepts | 0.709±0.126 | 0.847±0.139 | 0.648±0.162 | 0.835±0.157 | 0.714±0.109 |
| **CLEAR-HPV (AW-*h*)** | 0.784±0.141 | 0.843±0.117 | 0.797±0.197 | 0.623±0.284 | 0.715±0.206 |
| **CLEAR-HPV (raw-*h*)** | 0.749±0.119 | 0.889±0.078 | 0.770±0.181 | 0.633±0.204 | 0.684±0.146 |

**Table 3:** Original backbone performance vs. CLEAR-HPV concept discovery across MIL backbones (on TCGA-HNSCC). For each attention-based MIL backbone, we report the original backbone performance and the corresponding results obtained by applying CLEAR-HPV for concept discovery on the latent h-space. Feat. Dim. denotes the dimensionality of the slide-level representation used for classification, and Interpret. indicates whether the representation yields explicit, human-interpretable morphologic concepts. Across all backbones, CLEAR-HPV achieves strong performance comparable to the original models. For MHMIL, performance metrics for individual attention heads and alternative head-aggregation strategies are provided in in Table S4.

| Backbone | Method | Feat. Dim. | Interpret. | ACC | AUC | Prec | Rec | F1 |
| --- | --- | --- | --- | --- | --- | --- | --- | --- |
| *Original backbone performance (no concept discovery)* |  |  |  |  |  |  |  |  |
| ABMIL | original | 1536 | ✗ | 0.818±0.051 | 0.879±0.062 | 0.867±0.098 | 0.725±0.104 | 0.765±0.068 |
| TransMIL | original | 1536 | ✗ | 0.784±0.057 | 0.899±0.052 | 0.800±0.098 | 0.698±0.085 | 0.734±0.072 |
| MHMIL | original | 1536 | ✗ | 0.791±0.079 | 0.879±0.068 | 0.825±0.134 | 0.685±0.160 | 0.723±0.116 |
| *CLEAR-HPV concept discovery* |  |  |  |  |  |  |  |  |
| ABMIL | AW-*h* | 10 | ✓ | 0.799±0.087 | 0.859±0.081 | 0.769±0.115 | 0.767±0.222 | 0.751±0.133 |
| ABMIL | raw-*h* | 10 | ✓ | 0.810±0.068 | 0.868±0.086 | 0.811±0.145 | 0.763±0.169 | 0.769±0.102 |
| TransMIL | AW-*h* | 10 | ✓ | 0.765±0.077 | 0.863±0.089 | 0.738±0.071 | 0.703±0.179 | 0.713±0.111 |
| TransMIL | raw-*h* | 10 | ✓ | 0.784±0.089 | 0.841±0.084 | 0.803±0.149 | 0.678±0.147 | 0.727±0.118 |
| MHMIL | AW-*h* | 10 | ✓ | 0.782±0.145 | 0.859±0.144 | 0.813±0.229 | 0.669±0.242 | 0.716±0.204 |
| MHMIL | raw-*h* | 10 | ✓ | 0.794±0.100 | 0.879±0.129 | 0.859±0.140 | 0.669±0.273 | 0.713±0.178 |

Baselines that do not operate in the *h*-space performed substantially worse. We find that heatmap-based grouping produced diffuse partitions with limited HPV separation, indicating that spatial saliency alone is insufficient for isolating meaningful morphologic patterns; the Dirichlet model emphasizes either global regularity or high flexibility, which can conflict with the localized, heterogeneous, and uneven distribution of HPV-associated morphologic patterns in histopathologic tissue, making these distributions difficult to fit reliably under weak slide-level supervision; encoder-based clustering achieved reasonable quantitative performance but showed weaker class-coherent organization than *h*-space-based concept discovery, both qualitatively and quantitatively. In particular, the added class-dominance analyses in Figure 3(E) and (F) show that encoder-space concepts exhibit substantially lower dominant-cluster purity and mass concentration than CLEAR-HPV concepts, indicating greater mixing of class-relevant morphology within clusters. Both metrics are defined in Supplemental Methods S1.

#### Recovery score: How well CLEAR-HPV retains predictive performance

To further assess how well the discovered concepts retain the predictive performance of the backbone MIL model, we also compute a “recovery score” that quantifies how much predictive performance (e.g., accuracy and AUC) can be recovered using only the discovered concepts from different methods (including our CLEAR-HPV). (Details on how to compute the recovery score are included in the Methods section.) Figure 2 shows the results. CLEAR-HPV (AW-*h*) and CLEAR-HPV (raw-*h*) achieve the highest recovery scores, demonstrating that attention-weighted latent structure supports faithful, interpretable concept decompositions. Baselines such as heatmap-based grouping and Dirichlet clustering achieve much lower recovery scores, indicating a limited ability to preserve the original model’s predictive performance.

**Figure 2:**
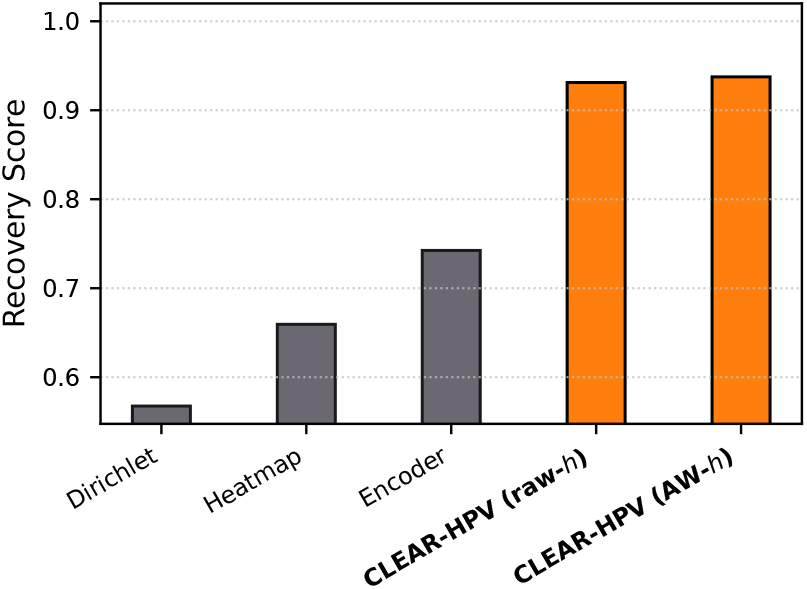
Recovery score relative to the interpreted MIL model (CLAM) across ACC, AUC, F1, Precision, Recall (i.e., sensitivity), and Specificity. For each method, the Euclidean distance *d* between its metric vector (i.e., concatenation of Accuracy, AUC, etc.) and the interpreted model’s is computed and converted to a similarity score 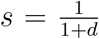. Higher scores indicate closer agreement with CLAM.

Together, these results show that concept discovery is most effective when performed directly in the attention-structured *h*-space, and that CLEAR-HPV can provide high-quality explanations of an MIL backbone’s prediction by retaining its discriminative signal while providing compact, interpretable concept-based representations. These are strong results because they show that CLEAR-HPV (1) preserves the predictive performance of the interpreted model while reducing its high-dimensional features (e.g., 1536 dimensions) to only 10 interpretable concepts and (2) improves class-coherent organization of discovered concepts, as quantified by the our class-dominance metrics.

### Interpretability

While the preceding section evaluated how well different concept-discovery methods preserve predictive signal and performance using quantitative classification metrics, these metrics alone do not explain what morphologic patterns the models rely on. We therefore next examine the interpretability of the discovered concepts by analyzing how they are distributed across slides and how clinicians can use these concept-fraction vectors. This analysis shifts the focus from performance to biological meaning, asking whether the learned concepts correspond to coherent and clinically recognizable histopathologic patterns.

#### Class-averaged concept-fraction analysis

To facilitate interpretability (explainability), we analyze concept-fraction vectors at the cohort level by averaging slide-level concept-fraction vectors within each group (e.g., HPV-positive and HPV-negative patients). These class-averaged concept-fraction vectors provide a compact quantitative summary of morphologic composition that can be directly compared across HPV status, survival outcomes, and cohorts. For example, an average concept-fraction vector over the HPV-positive patients provide insight on what concepts are most relevant to the positive predictions, and similarly for HPV-negative patients. A larger difference between these two vectors indicates that the discovered CLEAR-HPV concepts better distinguish HPV-positive from HPV-negative cases.

Figure 3 shows the class-averaged concept-fraction vectors for HPV-positive and HPV-negative cases with *K* = 10 concepts (C0–C9). Encoder-space concepts (Figure 3A) show little separation between classes. For CLEAR-HPV concepts discovered in the MIL *h*-space, unweighted concept fractions (Figure 3B; each tile contributes equally to its assigned concept) yield only modest class separation. In contrast, attention-weighted concept fractions (Figure 3C; each tile’s contribution to the slide-level fraction is weighted by its MIL attention score) produce a clearer dichotomy: HPV-positive slides show higher fractions of the basaloid/non-keratinizing concept (C5), whereas HPV-negative slides show higher fractions of the keratinizing concept (C7). Representative tiles supporting these morphologic interpretations are shown in Figure 4, consistent with established histopathologic differences. Using the same attention-weighted fraction computation, Figure 3D shows a clear survival-associated discrepancy in concept composition. Survivors are enriched for C5, whereas deceased cases are enriched for C7. This pattern is consistent with the established prognostic advantage of HPV-driven tumors, even though the MIL backbone was trained only for HPV prediction.

**Figure 3:**
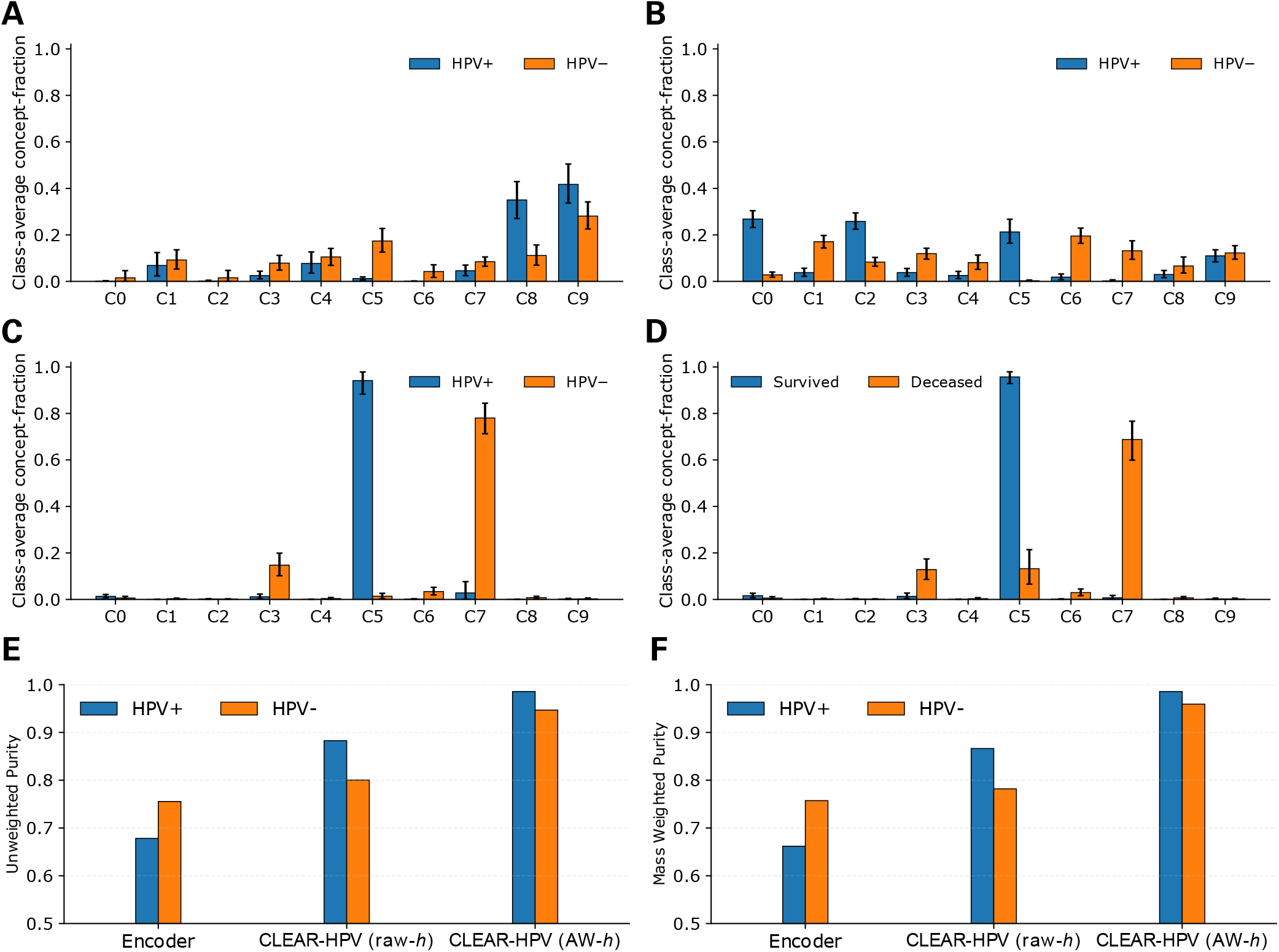
Class-averaged concept-fraction vectors across concept-discovery settings on TCGA-HNSCC. Concept-fraction vectors are computed per slide as the fraction of tiles assigned to each discovered concept, optionally weighted by MIL attention. These slide-level vectors are then averaged within each group to obtain class-averaged profiles that summarize cohort-level morphologic composition and highlight differences in concept prevalence across clinical groups. Panels (A–C) show group-averaged profiles for HPV-positive (blue) and HPV-negative (orange) cases, while panel (D) shows the corresponding averages for surviving (blue) and deceased (orange) cases. **(A)** Class-averaged concept-fraction vectors derived from encoder-feature clustering (non-*h*-space baseline). **(B)** Class-averaged concept-fraction vectors derived from CLEAR-HPV concepts in the MIL *h*-space using unweighted fractions, where all tiles contribute equally. **(C)** Class-averaged concept-fraction vectors derived from CLEAR-HPV concepts in the MIL *h*-space using attention-weighted fractions, where each tile contributes proportionally to its MIL attention score, yielding clearer separation between HPV-positive and HPV-negative cases. **(D)** Using the same CLEAR-HPV *h*-space concepts and attention-weighted fractions as in (C), class-averaged concept-fraction vectors are shown for surviving versus deceased cases. **(E)** Unweighted purity (cluster-averaged class dominance) across methods, computed as the mean class-dominance ratio over clusters dominated by each class. **(F)** Mass weighted purity (dominant-cluster class-mass concentration) across methods, computed as the fraction of class-specific mass within clusters dominated by each class; higher values indicate stronger class coherence. Detailed definitions are provided in Supplemental Methods S1.

**Figure 4:**
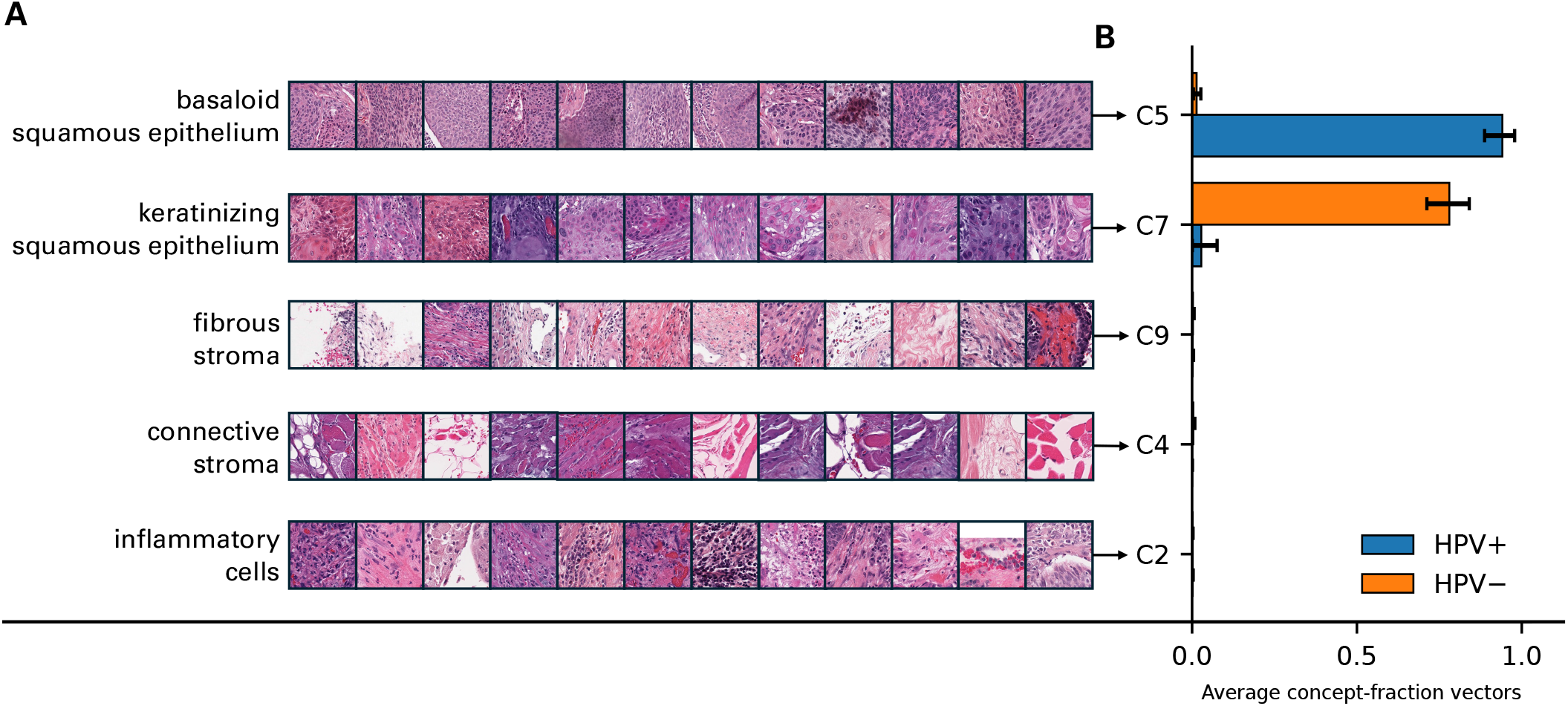
Top tiles for key concepts discovered by CLEAR-HPV (A) and the corresponding slide-level distributions in the dataset TCGA-HNSCC (B). **(A)** Top (representative) tiles for five CLEAR-HPV concepts chosen for their consistent appearance and clear morphologic identity: C5 (basaloid squamous epithelium), C7 (keratinizing squamous epithelium), C9 (fibrous stroma), C4 (connective stroma), and C2 (inflammatory cells). **(B)** Average concept-fraction vectors for HPV-positive (blue) and HPV-negative (orange) slides, with 95% bootstrap confidence intervals. The “Basaloid” Concept C5 is more prevalent in HPV-positive cases, while the “keratinizing” Concept C7 is more prevalent in HPV-negative cases.

To quantify class-coherent structure of discovered concepts, we introduce two metrics based on class-averaged concept-fraction vectors: unweighted purity, which measures the mean purity of class-dominant clusters, and mass weighted purity, which emphasizes whether class-specific signal is concentrated in high-mass clusters. Detailed definitions are provided in Supplemental Methods S1. Both metrics consistently show that attention-weighted CLEAR-HPV produces higher class-dominance and mass concentration than encoder-space concepts, indicating improved class-coherent organization of morphology.

#### Concept identity and representative morphology

Figure 4 shows representative tiles for key concepts discovered by CLEAR-HPV (Figure 4(A) and the corresponding slide-level distributions in TCGA-HNSCC (Figure 4(B)). In Figure 4(A), clinician review confirmed that these concepts correspond to inflammatory infiltrates (C2), benign stroma (C4), HPV-associated basaloid carcinoma (C5), HPV-negative keratinizing carcinoma (C7), and fibrous stroma (C9). These concepts appear consistently across datasets and form the basis for interpretable (explainable) slide-level composition profiles. In Figure 4(B), we can see that the discovered Concept 5 (C5) and Concept 7 (C7) are highly relevant to HPV-positive and HPV-negative cases, respectively.

#### Slide-level visualization of concepts discovered by CLEAR-HPV

Figure 5 visualizes concepts within two representative HPV-positive and HPV-negative slides. High-attention maps (Column 3 of Figure 5(A)) are obtained by ranking tile-level attention scores and retaining only top-scoring tiles, capturing regions most emphasized by the MIL backbone. Within these areas, HPV-positive slides consistently show the “basaloid” concept C5 (in yellow), while HPV-negative slides show the “keratinizing” concept C7. Background concepts such as fibrous or benign stromal tissue (e.g., C4 and C9), as indicated by their representative tiles in Figure 4, also appear consistently across slides and provide visual context for surrounding tumor regions. These representative tiles are selected by their distance to concept center (see details in the Methods section).

**Figure 5:**
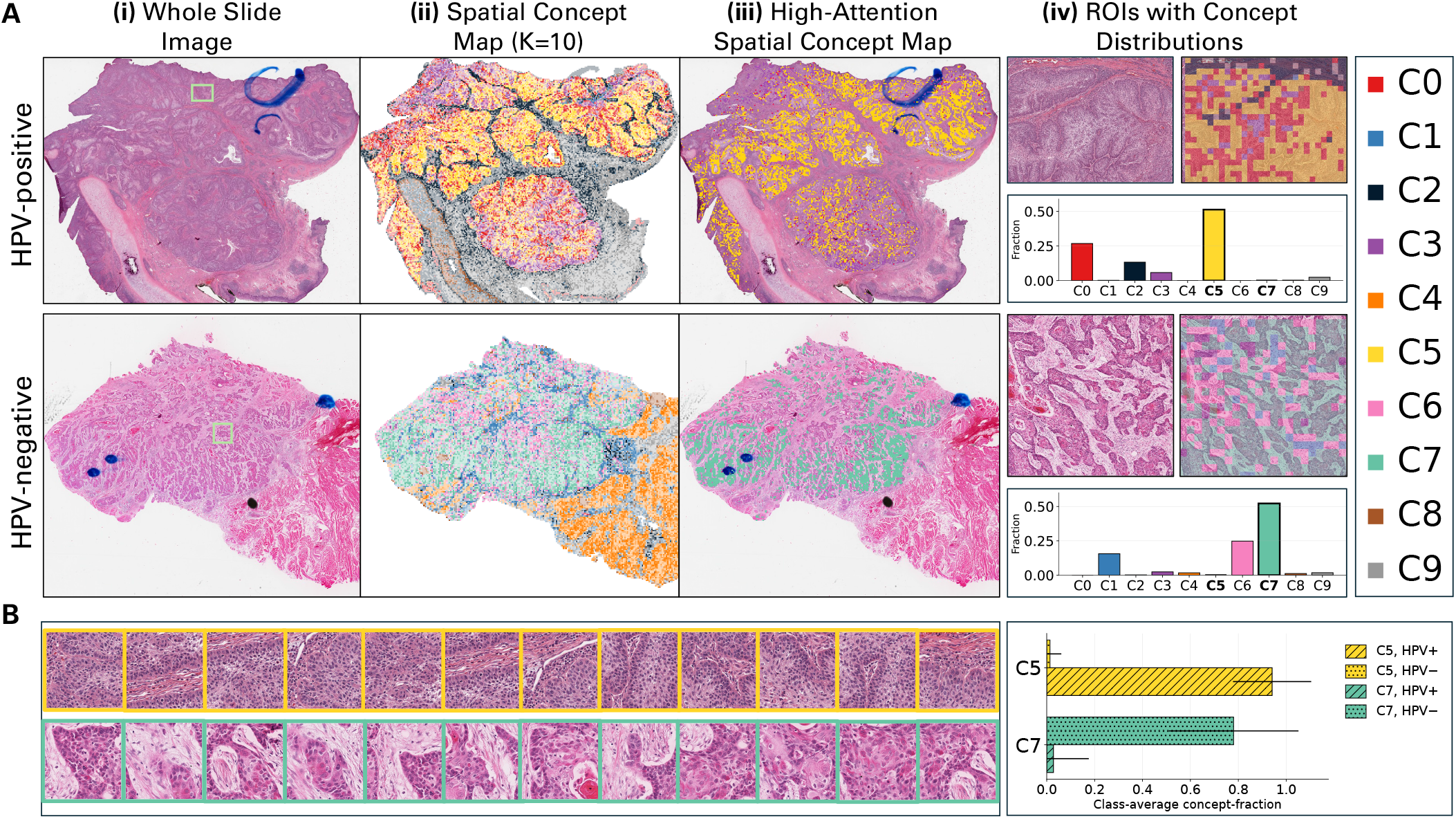
Visualization of attention-weighted concept discovery using CLEAR-HPV. **(A)** For representative HPV-positive and HPV-negative WSIs from TCGA-HNSCC, we show, in **four columns**: (***i***) the original H&E whole slide image, (***ii***) the *h*-space spatial concept map, (***iii***) the high-attention spatial concept map, and (***iv***) regions of interest (ROIs) with their corresponding concept-fraction distributions produced by our CLEAR-HPV. **(B)** Representative tiles for two CLEAR-HPV concepts (C5 and C7), along with the average concept-fraction vectors for HPV-positive and HPV-negative slides of the entire dataset. We use different colors to represent different concepts consistently across all figures. Blue markings visible on the whole slide images in (A) are pre-existing annotation artifacts on physical slides by clinicians and can be omitted from interpretation.

To quantify concept composition in highlighted, key areas of a slide, we define regions of interest (ROIs) as the contiguous high-attention regions outlined by the green box in Figure 5A(i), with a zoomed view shown in Figure 5A(iv) The concept-fraction distributions in the regions of interest (ROIs), shown in the green box in Figure 5(A)(i) and in Figure 5(A)(iv), match the dataset-level average concept-fraction vectors in Figure 5(B)(right): C5 is enriched in HPV-positive cases, and C7 is enriched in HPV-negative cases. Together, these results show that (1) our CLEAR-HPV can provide slide-level concept explanations for HPV prediction while also highlighting important regions in the slide, and that (2) our CLEAR-HPV produces consistent concepts at the slide and dataset levels; slide-level average concept-fraction vectors within high-attention regions align with their dataset-level (global) average concept-fraction vectors across the cohort.

### Comparison of concept discovery across MIL backbones

Our CLEAR-HPV framework is compatible with arbitrary attention-based MIL neural network architectures. In addition to CLAM, we evaluated three MIL backbones, including two widely used MIL backbones, i.e., attention-based MIL (ABMIL)^15^ and Transformer-based MIL (Trans-MIL)^17^, as well as a multi-head attention-based MIL model (MHMIL), to investigate how backbone choices influence the structure of the latent *h*-space and the resulting CLEAR-HPV concepts.

ABMIL employs a gated attention mechanism to learn instance-level importance weights and aggregates tile embeddings via a weighted sum, providing a simple and effective attention-based pooling strategy. TransMIL extends the MIL paradigm by modeling global contextual relationships among instances using Transformer layers, enabling long-range dependency modeling across tiles within a WSI. MHMIL is a multi-head attention-based MIL model inspired by CLAM; it replaces the single attention head with multiple parallel heads, allowing different heads to capture complementary tissue patterns while preserving the underlying MIL aggregation structure. By applying CLEAR-HPV to ABMIL, TransMIL, and MHMIL, we assess its robustness of concept discovery across distinct architectural inductive biases.

Using the same evaluation procedure as in the CLAM-based experiments, each slide is represented by a concept-fraction vector and evaluated with a concept-fraction classifier across all backbones. As shown in Table 3, CLEAR-HPV consistently preserves slide-level predictive performance across multiple attention-based MIL architectures, even though it replaces the original *high-dimensional* latent representations (i.e., 1536 dimensions), which are *not* interpretable at the concept level, with substantially more *compact* (i.e., 10 dimensions) and *interpretable* concept representations. These are therefore *strong* results. Across ABMIL, TransMIL, and MHMIL, CLEAR-HPV achieves AUC (≈0.84–0.88) and accuracy (≈0.76–0.81) that closely match those of the corresponding original backbone models (AUC ≈ 0.88–0.90, ACC ≈ 0.78–0.82). Importantly, we observe the following:

- Performance retention is consistent across architectures with distinct attention mechanisms and modeling complexity, indicating that the discovered concepts capture backbone-agnostic, diagnostically relevant morphology rather than architecture-specific artifacts.
- High performance is achieved when compressing the 1536-dimensional representation from the MIL backbone to only 10 interpretable concept dimensions, demonstrating that CLEAR-HPV retains discriminative signal in a substantially more compact and interpretable representation, without reliance on high-dimensional, uninterpretable embeddings.

#### Encoder dependence

In addition to backbone robustness, we evaluated whether CLEAR-HPV depends on the choice of feature encoder by replacing the UNI encoder with a conventional ResNet50. CLEAR-HPV maintains comparable performance under this change and continues to produce coherent concept representations, indicating that the proposed framework is not tied to a specific foundation model encoder. Detailed results are provided in Supplementary Table S5.

### Cross-cohort generalization and robustness

CLEAR-HPV concepts transfer reliably across datasets that differ in anatomic sites, staining profiles, and class distributions. All cross-cohort experiments were conducted in a zero-shot setting: concept clusters, attention weights, and the concept-fraction classifier were trained exclusively on TCGA-HNSCC and were not adapted, recalibrated, or tuned using any slides from TCGA-CESC or CPTAC-HNSCC. This design ensures that evaluations on the external cohorts measure genuine generalization rather than domain-specific adjustment. These results should be interpreted in terms of signal preservation and interpretability, rather than improvement in predictive performance.

#### Transferring to the external cohort TCGA-CESC

Table 4 shows the results when models trained on TCGA-HNSCC were applied to the external cohort TCGA-CESC. Both CLEAR-HPV variants, AW-*h* and raw-*h*, retain substantial predictive signal under this substantial biological shift in tissue site. Note that TCGA-CESC differs substantially from TCGA-HNSCC in both anatomy and cohort composition: it is dominated by HPV-positive tumors and exhibits histomorphologic patterns distinct from HNSCC^29^. Under this extreme imbalance, the CLAM baseline achieved only modest performance (ACC ≈ 0.59, AUC ≈ 0.68). Due to the strong class imbalance in TCGA-CESC, threshold-based metrics such as F1 and Recall should be interpreted with caution, as they can be influenced by majority-class bias. Interestingly, both CLEAR-HPV variants, with their discovered interpretable concepts and concept-fraction vectors, maintain comparable AUC while achieving higher accuracy (ACC ≈ 0.75), and, more importantly, preserved CLAM’s extremely high precision (Prec ≈ 0.96); they also achieved higher sensitivity and F1 than the CLAM base model. The attention-weighted (AW) variant achieved the highest F1 (≈0.86) and Sensitivity (≈0.78), while the raw-*h*-space variant obtained the highest AUC (≈0.67) among CLEAR-HPV variants. However, AUC indicates a modest degradation in ranking performance under domain shift, consistent with our interpretation that CLEAR-HPV preserves interpretable structure rather than improving global discrimination. These results show that CLEAR-HPV captures HPV-related morphologic signals that remain predictive, while maintaining interpretability under major shifts in tissue architecture. Together with the cross-cohort concept consistency shown in Fig. 6, these results indicate that CLEAR-HPV preserves transferable morphologic structure even when ranking performance degrades modestly.

**Table 4:**
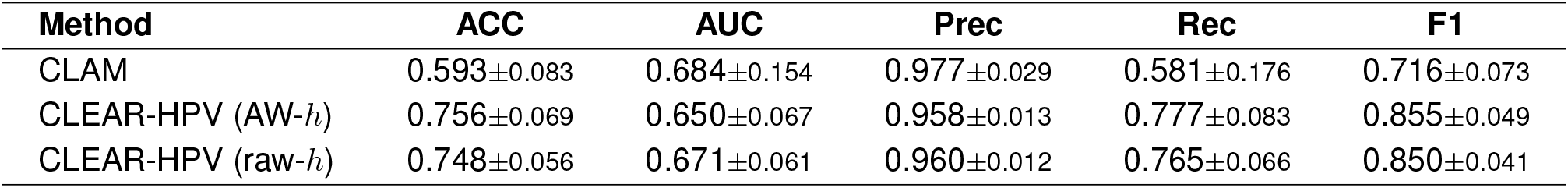
Cross-cohort generalization from TCGA-HNSCC to TCGA-CESC. Under severe domain shift and strong HPV-positive class imbalance, CLEAR-HPV retains substantial predictive signal in an interpretable concept space, although AUC indicates modest degradation in ranking performance relative to CLAM. raw-*h* achieves the highest AUC among the CLEAR-HPV variants, while AW-*h* provides the highest F1 and Recall. Results for additional metrics are provided in Table S6

**Figure 6:**
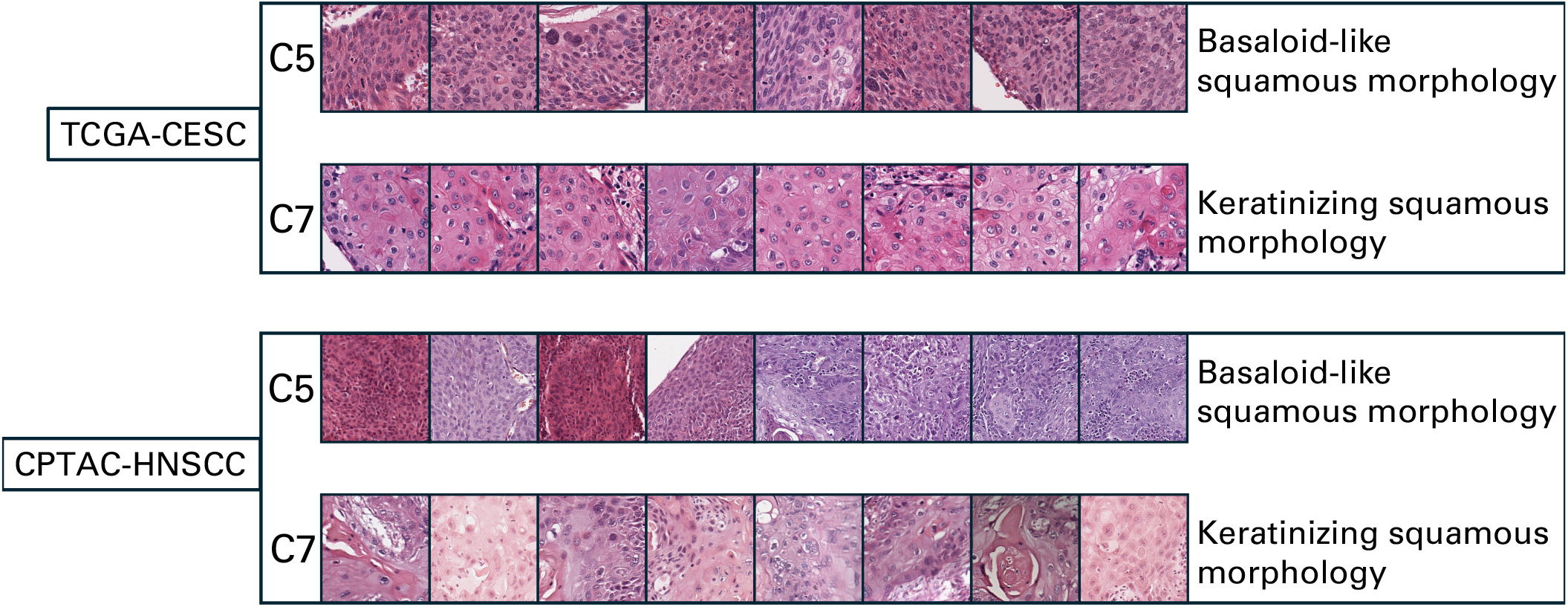
Cross-cohort consistency of HPV-related concepts among top-8 tiles. We show representative high-attention tiles for the HPV-positive-related “basaloid” concept C5 and the “keratinizing” concept C7 from two external cohorts, TCGA-CESC (**top**) and CPTAC-HNSCC (**bottom**). Across both datasets, C5 consistently reflects basaloid morphology characteristic of HPV-positive tumors, while C7 reflects keratinizing morphology typical of HPV-negative tumors. The consistency of these signatures across independent cohorts demonstrates that CLEAR-HPV identifies stable, biologically meaningful concepts that generalize beyond the training dataset.

#### Transferring to the external cohort CPTAC-HNSCC

Similarly, Table 5 shows the results when models trained on TCGA-HNSCC were applied to the external cohort CPTAC-HNSCC, demonstrating CLEAR-HPV’s robustness to technical and institutional variability. Because this cohort contains only HPV-negative tumors, accuracy (ACC) is the only relevant metric. Both CLEAR-HPV variants outperformed CLAM (ACC ≈ 0.82–0.90), despite pronounced differences in staining, slide preparation, and scanner characteristics^28^ between these two cohorts. This improvement indicates that our CLEAR-HPV’s concept-fraction representations (vectors) remain stable under technical domain shift.

**Table 5:** External validation on CPTAC-HNSCC. Only accuracy is reported because the cohort contains only HPV-negative cases. CLEAR-HPV preserves the predictive information of the CLAM base model when transferred to an external cohort.

| Metric | CLAM | CLEAR-HPV (AW-*h*) | CLEAR-HPV (raw-*h*) |
| --- | --- | --- | --- |
| ACC | 0.704±0.120 | 0.824±0.146 | 0.896±0.094 |

#### Morphologic consistency of concepts across cohorts

Figure 6 illustrates that the HPV-associated “basaloid” concept (C5) and the “keratinizing” concept (C7) retain their characteristic morphology in high-attention regions across both external cohorts, TCGA-CESC and CPTAC-HNSCC. The same concepts, basaloid tiles for C5 and keratinizing squamous morphology for C7, appear consistently even in these external cohorts not seen during training. These cross-cohort results demonstrate that CLEAR-HPV captures stable, biologically meaningful morphologic structures that generalize across different squamous tumor types and institutional settings.

### Survival prediction based on CLEAR-HPV concepts

HPV status is a well-established prognostic marker in squamous cell carcinoma, with HPV-positive tumors exhibiting significantly better survival outcomes^30^. We therefore assessed whether CLEAR-HPV concepts retain this prognostic signal by evaluating survival prediction in TCGA-HNSCC using only the concept-fraction vectors derived using CLEAR-HPV applied to an HPV-trained backbone. No retraining or survival-specific adaptation was performed; therefore the survival model receives solely the morphologic composition learned from HPV supervision.

As shown in Table 6, the attention-weighted concept vectors produced by CLEAR-HPV achieve competitive, though slightly lower, predictive performance compared to the base model CLAM. While the explained model, CLAM, achieves the highest AUC (≈ 0.840), both CLEAR-HPV variants (which explain CLAM’s predictions) retain broadly comparable discriminative ability despite operating in a substantially lower-dimensional space. Note that this is already strong results because the CLEAR-HPV’s concept-fraction vectors have only 10 dimensions; they are both compact and interpretable. In contrast, the original CLAM model uses embeddings with 1536 dimensions, and are not interpretable.

**Table 6:** Survival prediction on TCGA-HNSCC. Survival prediction using CLEAR-HPV concept representations (i.e., concept-fraction vectors) is compared with the base model CLAM. Results show that CLEAR-HPV’s concept-fraction vectors preserve the prognostic information contained in the original MIL embeddings of CLAM. Results for more metrics are provided in Table S7.

| Method | ACC | AUC | Prec | Rec | F1 |
| --- | --- | --- | --- | --- | --- |
| CLAM | 0.734±0.086 | 0.840±0.081 | 0.668±0.160 | 0.673±0.188 | 0.628±0.143 |
| CLEAR-HPV (AW-*h*) | 0.715±0.121 | 0.785±0.134 | 0.665±0.209 | 0.558±0.147 | 0.594±0.168 |
| CLEAR-HPV (raw-*h*) | 0.700±0.139 | 0.766±0.114 | 0.675±0.229 | 0.530±0.198 | 0.563±0.192 |

These results indicate that CLEAR-HPV concepts capture broader morphologic structure that remains informative for patient outcome, even though the concepts were originally learned for HPV prediction. The concept-fraction vectors provide a compact and interpretable summary of tumor and stromal composition, allowing links between morphologic patterns and adverse prognosis to be examined. Together, these findings show that reorganizing the MIL latent space into CLEAR-HPV concepts preserves survival-relevant information, even when supervision is based entirely on HPV labels.

## DISCUSSION

This study demonstrates that the latent space learned by attention-based MIL models contains biologically structured information that becomes interpretable when it is reorganized through our proposed CLEAR-HPV framework. The latent *h*-space encodes tile-level morphology before slide-level aggregation. Weighting these embeddings by learned attention scores highlights the regions most relevant to the prediction and enables CLEAR-HPV to recover consistent and generalizable morphologic concepts without tile-level supervision or human annotation. CLEAR-HPV is not designed to improve classification accuracy, but to provide a structured and interpretable view of the latent space learned by the model.

Our findings further demonstrate that CLEAR-HPV is not tied to a specific attention mechanism or MIL architecture, but can be applied consistently across diverse attention-based backbones. Despite differences in model architecture and attention formulation, CLEAR-HPV relies on the attention learned for the prediction task to organize the latent *h*-space, resulting in stable and robust morphologic concept discovery. This architectural robustness indicates that CLEAR-HPV functions as a general concept discovery framework that can be integrated with a wide range of attention-based MIL models without requiring model-specific modification.

CLEAR-HPV concepts demonstrate cross-cohort consistency across TCGA-HNSCC, TCGA-CESC, and CPTAC-HNSCC in a strict zero-shot transfer setting. Despite substantial variation in staining, scanner characteristics, and cohort composition^31,32^, the same “basaloid” and “keratinizing” concepts appeared consistently across cohorts (datasets). This reproducibility and stability indicate that CLEAR-HPV captures transferable morphologic signals associated with HPV status rather than dataset-specific artifacts, even under substantial domain shift, although some degradation in ranking performance is observed in certain transfer settings. Importantly, these results reinforce that CLEAR-HPV is designed to preserve predictive signal in an interpretable representation rather than to optimize classification performance, and thus minor changes in standard metrics should be interpreted in the context of this trade-off.

The concept-fraction vectors produced by CLEAR-HPV also retained prognostic information (important for survival analysis) in TCGA-HNSCC, even though the concepts were learned solely from HPV supervision without any concept-level annotation or supervision. This shows that reorganizing the MIL latent space can preserve broader tumor biology and that interpretable concept profiles can serve as compact descriptors for downstream clinical tasks.

While recent progress in pathology foundation models (e.g., contrastive histology encoders^33^, TCv2^34^, UNI^35^, CONCH^36^, and GigaPath^37^) provides powerful feature extractors trained at scale, our results highlight that expressive embeddings alone do not guarantee interpretability (or explainability). Direct clustering of foundation-model features produced reasonable quantitative performance but did not reveal coherent morphologic structure. The attention-weighted *h*-space, by contrast, combines the representational richness of these encoders with task-specific focus, enabling concept discovery that aligns with the regions most relevant to the prediction. This intermediate representation offers a practical pathway toward interpretable foundation-model pipelines.

While CLEAR-HPV organizes latent representations into structured and quantitatively coherent concepts, assigning semantic labels to these concepts currently requires expert interpretation. Future work may explore the integration of large language models (LLMs) or multimodal vision-language models to assist in automatic concept description or naming, for example by summarizing representative tiles associated with each concept. Such approaches could improve scalability and usability, but any automatically generated annotations would still require expert validation for clinical use.

In summary, CLEAR-HPV provides an interpretable link between slide-level predictions and human-understandable histology by restructuring the attention-mediated latent space into discrete morphologic concepts. These concepts are biologically coherent, reproducible across cohorts, and informative for classification and prognosis. More broadly, our findings show that the attention-structured latent space can support concept-level interpretability in whole-slide imaging across diverse MIL backbone architectures, offering a general framework for interpretable MIL. We expect that similar strategies will extend to additional molecular and prognostic tasks and will contribute to interpretable representation learning in computational pathology.

### Limitations of the study

This study has several limitations. First, the quality of the recovered concepts is influenced by the choice of MIL backbones; models with diffuse and weaker attention may offer slightly weaker structure for unsupervised separation of different concepts. Second, clustering is performed in a latent space without tile-level labels; therefore certain distinctions may remain subtle and benefit from expert interpretation. In particular, clinically actionable interpretation of the discovered concepts requires expert pathology review. Third, our pipeline relies on pretrained encoders and fixed attention maps, following the standard histopathology MIL paradigm (e.g., CLAM), where tile encoders are not jointly optimized during slide-level training. A unified end-to-end optimization of the encoder, attention mechanism, and concept structure may yield more refined and semantically consistent concepts, and remains an important direction for future work. Future work will incorporate spatial context, richer supervisory signals, and multimodal information to expand the biological insight provided by the CLEAR-HPV framework.

## METHODS

Figure 1 shows the overview of our CLEAR-HPV framework. Whole slide images (WSIs) are first decomposed into fixed-size tiles, encoded with a pretrained ViT or CNN, and converted into patch-level feature embeddings (Figure 1(A)). An attention-based MIL classifier then projects embeddings into an *h*-space latent representation and uses multi-head attention to compute tile-level contributions, according to which tile-level embeddings are then pooled into a single slide-level embedding for HPV prediction (Figure 1(B)).

Given an arbitrary attention-based MIL backbone model above, our CLEAR-HPV then performs annotation-free concept discovery on attention-weighted *h*-space embeddings to identify coherent morphologic concepts (Figure 1(C)). Using the discovered concepts, CLEAR-HPV can:

- compute the class-averaged concept-fractions to quantify the relative abundance of each concept for HPV-positive and HPV-negative slides within a given cohort; using attention weights can produce concepts that better distinguish between HPV-positive and HPV-negative cases (Figure 1(D)),
- generate representative tiles that illustrate the characteristic morphology captured by each discovered concept (Figure 1(E)),
- generate spatial concept maps to visualize the distribution of concepts across the WSIs, revealing their spatial organization (Figure 1(F)), and
- obtain slide-level concept-fraction vectors to provide an interpretable representation that supports a concept-fraction classifier, which recovers MIL predictive performance while offering concept-level explanations (Figure 1(G)).

Below, we provide details on each aformentioned module, evaluation, and implementation.

### Cohort curation and pathology criteria

TCGA-HNSCC WSIs were curated by removing slides with poor staining, marked artifacts, or inadequate focus, using a combination of metadata filtering and manual quality inspection. For TCGA-CESC, we followed the pathology quality-control criteria described in TCGA^29^ and retained only diagnostic slides with adequate tumor content and acceptable image quality. CPTAC-HNSCC slides were included as provided, since this cohort had already underwent standardized acquisition and quality review through the CPTAC pipeline.

### Data and preprocessing

We used diagnostic WSIs from TCGA-HNSCC, TCGA-CESC, and CPTAC-HNSCC. Tissue regions were identified using standard color-based masking, and each WSI was tiled into non-overlapping patches (256 *×*256) at the full-resolution layer (level 0), as shown in Figure 1(A). For each tile, we extracted a fixed embedding 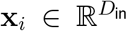 using the pretrained UNI encoder^35^ (*D*_in_ = 1536) without any color normalization. Each slide with *N* tiles is represented as an *N ×D*_in_ matrix of tile features. We have run experiments with color normalization too, but it turned out that color normalization does not make a difference, probably because the UNI encoder is already robust to different color schemes. Therefore we chose to run all experiments without any color normalization.

### MIL backbones and the *h*-space

#### Different MIL backbones

We evaluated four attention-based MIL architectures, including CLAM, ABMIL, TransMIL, and MHMIL. All models share a unified projection layer that maps encoder features into a common latent *h*-space with the same dimensionality, enabling direct and controlled comparison of the resulting representations across backbones, See Figure 1(B) for a typical example of attention-based MIL model.

#### Attention-structured *h*-space

As shown in Figure 1(B), we define the *h*-space (the blue box) as the intermediate embedding space produced by the MIL backbone prior to slide-level aggregation. For each tile *i*, the backbone maps the encoder feature **x**_*i*_ to an embedding **h**_*i*_ = *f*_*θ*_(**x**_*i*_) and produces a raw attention score *a*_*i*_ (i.e., the logit before “softmax”). Attention weights are obtained via slide-wise softmax over the *N* tiles in the slide, followed by a rescaling step:

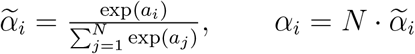

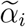 denotes the normalized attention weight, and *N* denotes the number of tiles extracted from the current slide. The rescaling sets the *mean* weight within each slide to 1 (since 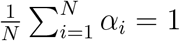), which makes the *average* magnitude of attention weights comparable across slides with different numbers of tiles while preserving the relative ranking of tiles within each slide. The **resulting attention-structured** *h***-space representation of a slide** is given by

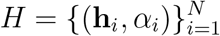

This standardized *h*-space representation above serves as the input to CLEAR-HPV’s unsupervised concept discovery procedure, which analyzes the attention-weighted distribution of tile embeddings to reveal structured morphologic patterns learned by the MIL model.

### Concept discovery module

CLEAR-HPV performs unsupervised concept discovery by clustering the attention-structured *h*-space representation into a fixed set of *K* latent morphologic concepts, *without requiring any concept-level annotations (labels)*. In Table 2, we compare CLEAR-HPV (AW-*h*) and CLEAR-HPV (raw-*h*) against several unsupervised baselines, including Heatmap, Dirichlet Concepts, and Encoder Concepts. We describe each method in detail below. All methods operate on tile-level *h*-space representation {(**h**_*i*_, *α*_*i*_)} and are designed to scale to whole-slide inference.

#### CLEAR-HPV (raw-*h*)

The raw variant of CLEAR-HPV, i.e., *CLEAR-HPV (raw-h)*, discovers *K* morphologic concepts by clustering tile embeddings in the *h*-space using a scalable two-stage k-means procedure^38,39^. Let 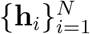 denote the *N* tile embeddings used for concept learning, where each 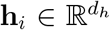 is a *d*_*h*_-dimensional feature vector for tile *i*. We learn *K* concept centroids 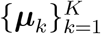 with 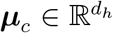 and a hard assignment function *c*(*i*) *∈* {1, …, *K*} that maps each tile *i* to its nearest centroid by minimizing the within-cluster sum of squared distances:

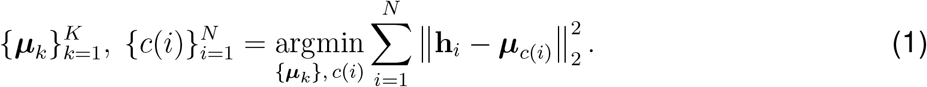

In practice, we initialize concept centroids in the *d*_*h*_-dimensional *h*-space using a reservoir-based mini-batch K-means pass, followed by standard K-means clustering^40^ applied to all tile embeddings to obtain stable concept centroids. This variant treats all tiles equally during concept learning. We next introduce *CLEAR-HPV (AW-h)*, which incorporates attention weights when forming concepts and provides a complementary view of the attention-structured *h*-space.

#### CLEAR-HPV (AW-*h*)

To incorporate the MIL model’s diagnostic relevance into concept learning, CLEAR-HPV (AW-*h*) uses an attention-weighted k-means objective. Let 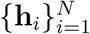 denote tile embeddings and let *α*_*i*_ ≥ 0 denote the corresponding attention weight assigned by the MIL model (with by solving ∑_*i*_ *α*_*i*_ *>* 0). We learn *K* centroids 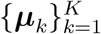 and hard assignments *c*(*i*) *∈ {*1, …, *K}* by solving

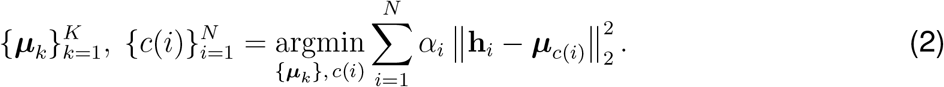

During initialization and refinement, tile weights *α*_*i*_ act as sample weights, causing high-attention tiles to exert stronger influence on concept boundaries. This aligns the learned concepts with regions emphasized by the MIL model.

#### Dirichlet concepts

Building on ideas from probabilistic concept models such as PACE^41^, we include a probabilistic concept model that learns uncertainty-aware importance score for each tile and each concept for the prediction. A probabilistic model like this is usually more robust and captures broader variability in morphologic patterns across slides.

#### Heatmap

We evaluated a heatmap-based slide classifier that uses the MIL attention map as a quantitative decision signal. For each slide, tile-level attention scores are projected back onto the WSI to form a spatial heatmap. A slide-level prediction is obtained by computing a scalar “heatmap area” score, defined as the proportion of tiles whose attention exceeds a fixed threshold *τ* for the class of interest. The decision cutoff is selected on training data and applied to held-out test slides, yielding a transparent rule-based classifier grounded in attention-based evidence.

#### Encoder concepts

As a baseline, we evaluate an encoder-based concept discovery method that follows the same procedure as CLEAR-HPV (raw-*h*) but operates directly on encoder feature space rather than the MIL latent *h*-space. Let 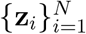, with 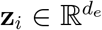, denote the tile-level embeddings produced by the pretrained encoder; all subsequent steps are identical to CLEAR-HPV (raw-*h*). Because this baseline does not involve MIL attention, an attention-weighted (AW-*h*) variant is not applicable.

#### Choice of number of concepts (*K*)

We use *K* = 10 as the main setting in our experiments. This choice was initially guided by elbow analysis^42^ and further supported by evaluations across nearby values of *K*. These analyses show that predictive performance remains stable and that the overall structure of the discovered concepts is highly consistent across different values of *K*. In practice, changing *K* mainly merges or splits similar concepts rather than producing entirely different patterns. We therefore do not interpret *K* = 10 as a biologically unique optimum, but as a stable and interpretable choice for this study.

### Concept-fraction Vectors in CLEAR-HPV

As shown in Figure 1(D) and Figure 1(G), our CLEAR-HPV provides both class-level and slide-level concept-fraction vectors to enable holistic interpretation (explanation) of MIL models’ predictions. Below we describe details on how they are computed.

#### Raw slide-level concept-fraction vectors

Given a set of *K* discovered concepts from CLEAR-HPV, we represent each whole slide image using a *concept-fraction vector* that summarizes its morphologic composition. Specifically, for a slide with *N* tiles, let *c*(*i*) *∈ {*1, …, *K}* denote the index of the concept assigned to tile *i ∈ {*1, 2, …, *N}*. The slide-level concept-fraction vector **f** *∈* ℝ^*K*^ is defined as

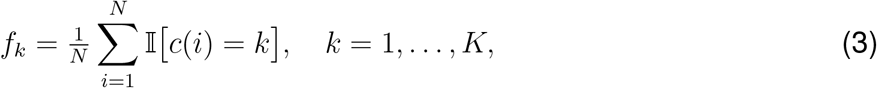

where *f*_*k*_ is the *k*-th entry in **f**, and 𝕀 [·] is the indicator function, equal to 1 if tile *i* is assigned to concept *k* and 0 otherwise. Each entry *f*_*k*_ therefore represents the proportion of tiles in the slide assigned to concept *k*, and the vector **f** is normalized such that 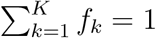.

#### Attention-weighted slide-level concept-fraction vectors

For attention-weighted variants, each tile *i* is additionally associated with an attention weight *α*_*i*_ produced by the MIL model, reflecting its contribution to the slide-level prediction. Correspondingly, the *attention-weighted (AW) concept fraction vectors* are computed as

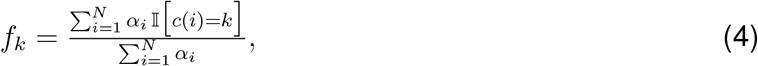

where tiles receiving higher attention contribute more strongly to the slide-level representation. Attention weights are normalized within each slide to ensure comparability across slides of different sizes.

These two variants of slide-level concept-fraction vectors are used as inputs to the concept-fraction classifier for quantitative evaluation.

#### Class-averaged concept-fraction vectors

For interpretability analyses, we further compute class-averaged concept-fraction vectors by averaging slide-level concept-fraction vectors across all slides within a given cohort or clinical group (e.g., averaging over HPV-positive cases or HPV-negative cases). Throughout the paper, we use the term concept-fraction vector to denote slide-level representations, and class-averaged concept-fraction vectors when these representations are averaged within a subgroup of a cohort (e.g., HPV-positive cases) for interpretability analyses.

### Representative tiles per concept

As shown in Figure 1(E), to support qualitative interpretation of the discovered concepts, we identify representative tiles for each concept using an automated and reproducible ranking procedure. Tiles are ranked by their Euclidean distance to the corresponding concept centroid 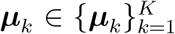, as defined by Eqn. (1), and the top *M* tiles are selected across slides for visualization. These representative tiles are used exclusively for qualitative interpretation and do not influence model training or evaluation.

### Spatial concept maps

#### Spatial concept maps

As shown in Figure 1(F) and Figure 5(A)(ii), to visualize the spatial organization of discovered concepts within WSIs, we generate spatial concept maps by projecting tile-level concept assignments back to their original spatial locations in the WSI. Each tile is assigned to a single concept using hard assignment, and tiles are colored according to their assigned concept, producing a map that illustrates how different morphologic concepts are distributed across the tissue section.

#### High-attention spatial concept maps

As shown in Figure 5(A)(iii), when attention scores are available, we additionally generate a high-attention map by displaying only the top *M* tiles ranked by MIL attention score, highlighting regions most strongly emphasized by the model.

### Evaluation of concept-fraction vectors

All concept-discovery methods were evaluated under a consistent 10-fold train/test protocol applied to the *h*-space. For each fold, concepts were learned using only training slides, and concept-fraction vectors were computed for both training and held-out slides. Slide-level predictions derived from concept fractions were compared against the baseline MIL classifier.

#### Concept-fraction classifier

Given a slide 𝒮, let **f** (𝒮) = [*f*_1_, …, *f*_*K*_] ∈ ℝ^*K*^ denote its slide-level concept-fraction vector as defined in Eqn. (3) and Eqn. (4), where *f*_*k*_ is the proportion of tiles assigned to concept *k* and 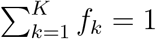. We then define a simple, interpretable rule-based classifier that scores each slide by aggregating the fractions of concepts that are statistically associated with HPV-positive status in the training data. Specifically, for each concept *k*, we compare the distribution of *f*_*k*_ between HPV-positive and HPV-negative training slides and designate concept *k* as HPV-associated if its mean fraction is higher in the HPV-positive group. This yields a binary concept mask ***π*** ∈ {0, 1} ^*K*^, where *π*_*k*_ = 1 indicates that concept *k* is positively associated with HPV status based on the training set, and *π*_*k*_ = 0 otherwise.

The slide-level score is computed as

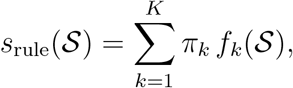

which represents the total fraction of tissue assigned to HPV-associated concepts. A binary prediction is obtained by thresholding this score,

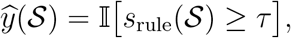

where 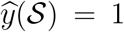 denotes an HPV-positive prediction and 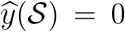 denotes an HPV-negative prediction. The decision threshold *τ* is selected using the training folds and then applied to held-out test slides.

For comparison, we also train a logistic regression classifier directly using the same concept-fraction vector **f** (𝒮) as input. Detailed performance metrics for HPV and survival prediction are reported in Supplementary Table S8 and Table S9.

#### Recovery score

Our CLEAR-HPV is a post-hoc explanation framework. It is therefore important to evaluate whether the generated explanation indeed reflects the explained model’s prediction. To measure how well CLEAR-HPV variants and other baselines recover MIL predictions, we computed a recovery score by forming a metric vector,

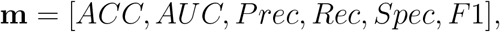

for each method (e.g., **m**_*CLEAR*−*HPV*_ for CLEAR-HPV and **m**_*Base*_ for a certain base model) and calculating its Euclidean distance *d* to the base model (e.g., CLAM) in the same fold. The final recovery score (for each fold) is

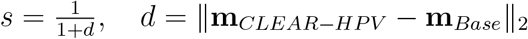

Supplemental Methods S1. Definition of class-dominance ratios and dominant-cluster metrics (Avg Ratio and Sum Ratio).

Figure S1. Elbow analysis used to select the number of morphologic concepts.

Table S1. Performance across different numbers of concepts (*K*) for CLEAR-HPV (AW-*h*) on TCGA-HNSCC.

Table S2. Stability of discovered concepts across different *K*, measured by forward persistence (FP) and reverse fragmentation (RF).

Table S3. Results in terms of complete evaluation metrics for concept discovery methods on TCGA-HNSCC.

Table S4. Effect of attention head selection in MHMIL-IR on TCGA-HNSCC.

Table S5. Encoder-dependence analysis using ResNet50 comparing encoder-space concepts and CLEAR-HPV.

Table S6. Results in terms of complete evaluation metrics for cross-cohort generalization from TCGA-HNSCC to TCGA-CESC.

Table S7. Results in terms of complete evaluation metrics for TCGA-HNSCC survival prediction using CLEAR-HPV concepts derived from an HPV-trained backbone.

Table S8. Logistic regression classifier-based HPV prediction on TCGA-HNSCC using CLEAR-HPV.

Table S9. Logistic regression classifier-based survival prediction on TCGA-HNSCC using CLEAR-HPV.

## Supporting information

all supplemental materials

