## Supplementary material for "CLEAR-HPV: Interpretable Concept Discovery for HPV-Associated Morphology in Whole-Slide Histology": all supplemental materials

### Supplemental Methods S1

**Class-dominance ratios.** Let $f_{i,c} \in [0,1]$ denote the concept-fraction of slide $i$ assigned to cluster (concept) $c$, where $c \in \{1,\ldots,K\}$ and $i$ indexes slides. Let $\mathrm{HPV}^+$ and $\mathrm{HPV}^-$ denote the sets of HPV-positive and HPV-negative slides, respectively, with sizes $N_+$ and $N_-$. The class-averaged concept fractions are:

$$\mu_c^+ = \frac{1}{N_+} \sum_{i \in \mathrm{HPV}^+} f_{i,c}, \qquad \mu_c^- = \frac{1}{N_-} \sum_{i \in \mathrm{HPV}^-} f_{i,c}. \tag{1}$$

We define normalized class-dominance ratios:

$$r_c^+ = \frac{\mu_c^+}{\mu_c^+ + \mu_c^- + \epsilon}, \qquad r_c^- = \frac{\mu_c^-}{\mu_c^+ + \mu_c^- + \epsilon}, \tag{2}$$

where $\epsilon > 0$ is a small constant for numerical stability. Values near 1 indicate strong class dominance, while values near 0.5 indicate mixed clusters.

**Dominant-cluster metrics: Unweighted purity and mass weighted purity.** Built on the class-dominance ratios above, we can then define our metrics. First, define dominant cluster sets:

$$\mathcal{D}_+ = \{c : \mu_c^+ > \mu_c^-\}, \qquad \mathcal{D}_- = \{c : \mu_c^- > \mu_c^+\}. \tag{3}$$

The *unweighted purity* is:

$$P_+^{\mathrm{unw}} = \frac{1}{|\mathcal{D}_+|} \sum_{c \in \mathcal{D}_+} r_c^+, \qquad P_-^{\mathrm{unw}} = \frac{1}{|\mathcal{D}_-|} \sum_{c \in \mathcal{D}_-} r_c^-. \tag{4}$$

The *mass weighted purity* is:

$$P_+^{\mathrm{mw}} = \frac{\sum_{c \in \mathcal{D}_+} \mu_c^+}{\sum_{c \in \mathcal{D}_+} (\mu_c^+ + \mu_c^-)}, \qquad P_-^{\mathrm{mw}} = \frac{\sum_{c \in \mathcal{D}_-} \mu_c^-}{\sum_{c \in \mathcal{D}_-} (\mu_c^+ + \mu_c^-)}. \tag{5}$$

Intuitively, these metrics assess whether each discovered concept is dominated by a single class (high coherence) or mixes multiple classes (low coherence), providing a quantitative proxy for alignment with class-specific morphology. Unweighted purity captures average cluster-level dominance, while mass weighted purity emphasizes the concentration of class-specific mass in dominant clusters; *higher values indicate better coherence.*

#### Supplemental Figures

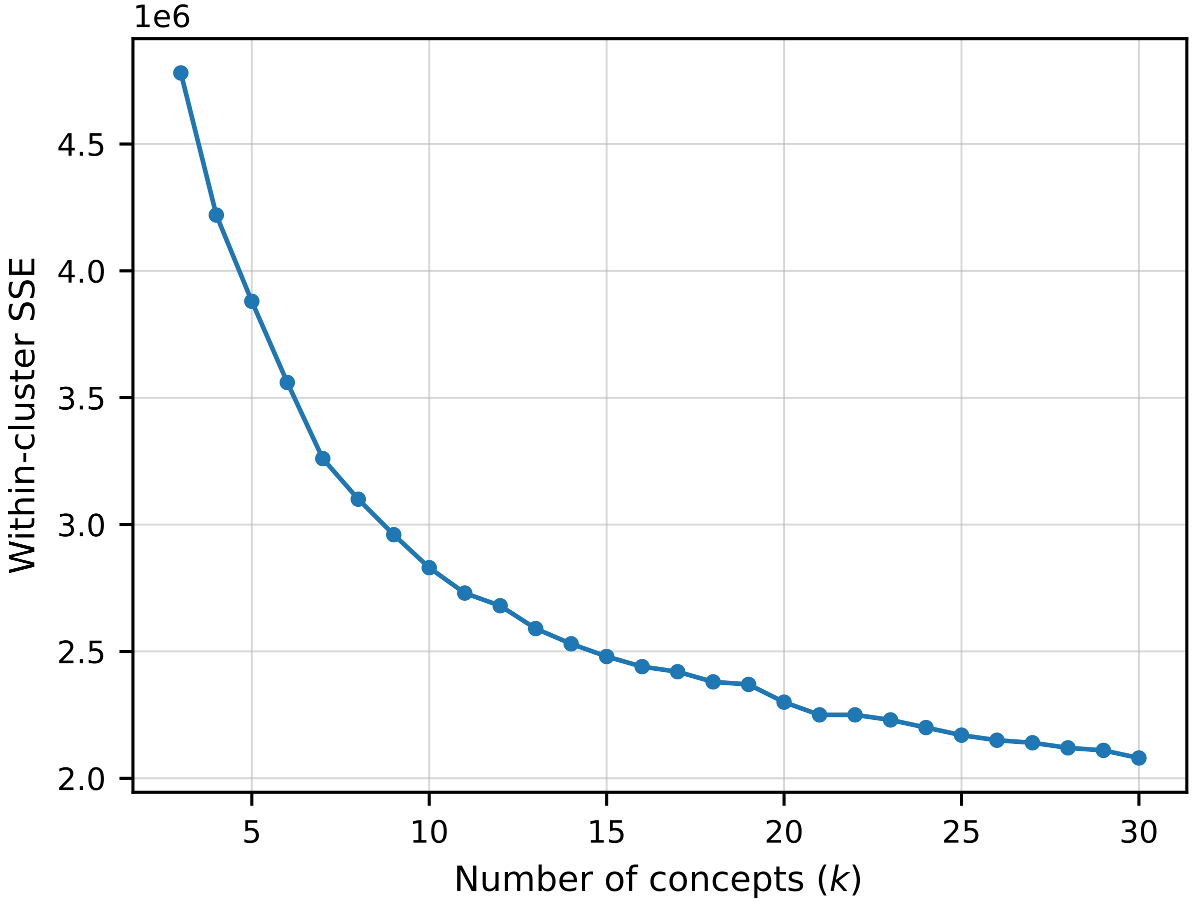


Figure S1: **"Elbow" analysis for concept number selection.** This "elbow" plot shows the within-cluster sum of squared errors (SSE) as a function of the number of concepts K for clustering tile embeddings in the CLAM $h$-space. The curve exhibits a diminishing rate of SSE reduction roughly beyond $K$=10, which was therefore selected as a balanced trade-off between compactness and clustering fidelity in CLEAR-HPV.

#### Supplemental Tables

| $K$ | ACC | F1 | Prec | Rec | Spec | AUROC |
|---|---|---|---|---|---|---|
| 10 | $0.773_{\pm 0.073}$ | $0.688_{\pm 0.121}$ | $0.815_{\pm 0.133}$ | $0.623_{\pm 0.153}$ | $0.883_{\pm 0.080}$ | $0.830_{\pm 0.081}$ |
| 5 | $0.757_{\pm 0.059}$ | $0.696_{\pm 0.072}$ | $0.788_{\pm 0.116}$ | $0.658_{\pm 0.122}$ | $0.833_{\pm 0.097}$ | $0.827_{\pm 0.069}$ |
| 15 | $0.747_{\pm 0.068}$ | $0.687_{\pm 0.086}$ | $0.753_{\pm 0.110}$ | $0.658_{\pm 0.122}$ | $0.817_{\pm 0.088}$ | $0.837_{\pm 0.063}$ |

Table S1: **Analysis across varying $K$ for CLEAR-HPV (AW-$h$) on TCGA-HNSCC.** Results remain stable across $K \in \{5, 10, 15\}$, with overlapping confidence intervals across all major metrics. This supports $K = 10$ as a balanced operating point rather than a uniquely optimized choice, providing comparable predictive performance while retaining the concept granularity used throughout the manuscript. Performance is reported as mean ± 95% confidence interval over 10 folds.

| $K$ | Forward Persistence (FP) | Reverse Fragmentation (RF) |
|---|---|---|
| 5 | $0.962_{\pm 0.012}$ | $0.988_{\pm 0.005}$ |
| 10 | – | – |
| 15 | $0.988_{\pm 0.002}$ | $0.982_{\pm 0.004}$ |

Table S2: **Centroid stability across different $K$ relative to $K = 10$.** Forward persistence (FP) measures how well concepts at $K = 10$ are preserved when compared against an alternative resolution, while reverse fragmentation (RF) measures how consistently concepts at the alternative resolution map back to the $K = 10$ reference. Values close to 1 indicate stable concept structure across resolutions. Values are reported as mean ± 95% confidence interval over 10 folds.

| **Method** | **ACC** | **AUC** | **F1** | **Prec** | **Rec** | **Spec** |
|---|---|---|---|---|---|---|
| Heatmap | $0.538_{\pm 0.110}$ | $0.674_{\pm 0.147}$ | $0.538_{\pm 0.114}$ | $0.503_{\pm 0.137}$ | $0.653_{\pm 0.180}$ | $0.486_{\pm 0.165}$ |
| Dirichlet Concepts | $0.646_{\pm 0.071}$ | $0.641_{\pm 0.134}$ | $0.266_{\pm 0.193}$ | $0.475_{\pm 0.314}$ | $0.200_{\pm 0.160}$ | $0.983_{\pm 0.033}$ |
| Encoder Concepts | $0.709_{\pm 0.078}$ | $0.847_{\pm 0.086}$ | $0.714_{\pm 0.068}$ | $0.648_{\pm 0.100}$ | $0.835_{\pm 0.096}$ | $0.617_{\pm 0.193}$ |
| CLEAR-HPV (AW-$h$) | $0.784_{\pm 0.087}$ | $0.843_{\pm 0.072}$ | $0.715_{\pm 0.128}$ | $0.797_{\pm 0.122}$ | $0.673_{\pm 0.148}$ | $0.867_{\pm 0.081}$ |
| CLEAR-HPV (raw $h$) | $0.749_{\pm 0.074}$ | $0.889_{\pm 0.049}$ | $0.684_{\pm 0.090}$ | $0.770_{\pm 0.112}$ | $0.633_{\pm 0.108}$ | $0.833_{\pm 0.084}$ |

Table S3: **Results in terms of complete evaluation metrics for concept discovery methods on TCGA-HNSCC.** Results are reported as accuracy (ACC), area under the ROC curve (AUC), F1, precision (Prec), recall (Rec, same as sensitivity), and specificity (Spec) as mean ± 95% confidence interval (CI) across cross-validation folds.

| Method | ACC | AUC | Prec | Rec | F1 |
|---|---|---|---|---|---|
| MHMIL-IR (head 0) | 0.804±0.105 | 0.866±0.103 | 0.857±0.150 | 0.669±0.173 | 0.737±0.149 |
| MHMIL-IR (head 1) | 0.802±0.101 | 0.848±0.101 | 0.857±0.150 | 0.659±0.172 | 0.731±0.146 |
| MHMIL-IR (head 2) | 0.782±0.104 | 0.855±0.104 | 0.820±0.159 | 0.641±0.176 | 0.706±0.150 |
| MHMIL-IR (head 3) | 0.770±0.108 | 0.852±0.105 | 0.798±0.173 | 0.637±0.152 | 0.700±0.149 |
| MHMIL-IR (max) | 0.772±0.099 | 0.852±0.105 | 0.809±0.164 | 0.646±0.154 | 0.704±0.139 |
| MHMIL-IR (mean) | 0.782±0.104 | 0.859±0.103 | 0.813±0.164 | 0.669±0.173 | 0.716±0.146 |
| MHMIL-IR (sum) | 0.782±0.104 | 0.859±0.103 | 0.813±0.164 | 0.669±0.173 | 0.716±0.146 |

Table S4: **Effect of attention head choices for MHMIL-IR on TCGA-HNSCC.** We report performance from CLEAR-HPV concept-fraction vectors when deriving attention-weighted $h$-space embeddings using individual MHMIL-IR attention heads (0–3) or simple head aggregations (max/mean/sum). Metrics are reported as mean ± 95% confidence interval across cross-validation folds.

| Method | ACC | AUC | Prec | Rec | Spec | F1 |
|---|---|---|---|---|---|---|
| ResNet50 + Encoder Concepts | 0.689±0.024 | 0.758±0.086 | 0.645±0.093 | 0.727±0.105 | 0.667±0.112 | 0.663±0.027 |
| ResNet50 + CLEAR-HPV (raw-$h$) | 0.700±0.066 | 0.800±0.083 | 0.670±0.104 | 0.637±0.104 | 0.750±0.084 | 0.643±0.086 |
| ResNet50 + CLEAR-HPV (AW-$h$) | 0.720±0.080 | 0.796±0.082 | 0.710±0.126 | 0.637±0.104 | 0.783±0.098 | 0.662±0.099 |

Table S5: **Encoder-dependence analysis with expanded metrics (ResNet50).** Comparison of encoder-space concept discovery (Encoder Concepts) and CLEAR-HPV in the ResNet50-based $h$-space under $K = 10$ and 10-fold evaluation. Using the same ResNet50 encoder features, CLEAR-HPV yields improved accuracy, AUC, precision, and specificity relative to encoder-space clustering, whereas recall decreases and F1 remains comparable. Metrics are reported as mean ± 95% confidence interval across cross-validation folds.

| Method | ACC | AUC | F1 | Prec | Rec | Spec |
|---|---|---|---|---|---|---|
| CLEAR-HPV (AW-$h$) | 0.536±0.055 | 0.702±0.046 | 0.674±0.053 | 0.977±0.011 | 0.520±0.060 | 0.800±0.102 |
| CLEAR-HPV (raw $h$) | 0.586±0.055 | 0.701±0.037 | 0.719±0.049 | 0.976±0.008 | 0.575±0.061 | 0.767±0.089 |

Table S6: **Results in terms of complete evaluation metrics for cross-cohort generalization from TCGA-HNSCC to TCGA-CESC.** Complete evaluation of cross-cohort transfer performance under a zero-shot setting, where all models are trained exclusively on TCGA-HNSCC and evaluated on TCGA-CESC without any target-cohort fine-tuning. Results are reported as accuracy (ACC), area under the ROC curve (AUC), F1, precision (Prec), recall (Rec, same as sensitivity), and specificity (Spec) as mean ± 95% confidence interval (CI) across cross-validation folds.

| Method | ACC | AUC | Prec | Rec | Spec | F1 |
|---|---|---|---|---|---|---|
| CLAM backbone | 0.734±0.086 | 0.840±0.081 | 0.668±0.160 | 0.673±0.188 | 0.778±0.096 | 0.628±0.143 |
| CLEAR-HPV (AW-$h$) | 0.715±0.121 | 0.785±0.134 | 0.665±0.209 | 0.558±0.147 | 0.812±0.128 | 0.594±0.168 |
| CLEAR-HPV (raw $h$) | 0.700±0.139 | 0.766±0.114 | 0.675±0.229 | 0.530±0.198 | 0.812±0.140 | 0.563±0.192 |

Table S7: **Results in terms of complete evaluation metrics for TCGA-HNSCC survival prediction using CLEAR-HPV concepts derived from an HPV-trained backbone.** The backbone is trained exclusively for HPV prediction, and CLEAR-HPV is then applied post hoc to this backbone. Results are reported as accuracy (ACC), area under the ROC curve (AUC), F1, precision (Prec), recall (Rec, same as sensitivity), and specificity (Spec) as mean ± 95% confidence interval (CI) across cross-validation folds.

| Method | ACC | AUC | Prec | Rec | Spec | F1 |
|---|---|---|---|---|---|---|
| CLEAR-HPV (AW-$h$) | 0.747$\pm$0.076 | 0.855$\pm$0.052 | 0.758$\pm$0.114 | 0.633$\pm$0.124 | 0.833$\pm$0.079 | 0.679$\pm$0.098 |
| CLEAR-HPV (raw $h$) | 0.731$\pm$0.086 | 0.851$\pm$0.059 | 0.765$\pm$0.129 | 0.592$\pm$0.151 | 0.833$\pm$0.097 | 0.644$\pm$0.122 |

Table S8: **Logistic regression classifier-based HPV prediction on TCGA-HNSCC using CLEAR-HPV.** Results are reported as accuracy (ACC), area under the ROC curve (AUC), F1, precision (Prec), recall (Rec, same as sensitivity), and specificity (Spec) as mean ± 95% confidence interval (CI) across cross-validation folds.

| Method | ACC | AUC | Prec | Rec | Spec | F1 |
|---|---|---|---|---|---|---|
| CLEAR-HPV (AW-$h$) | 0.735$\pm$0.116 | 0.827$\pm$0.105 | 0.720$\pm$0.201 | 0.588$\pm$0.211 | 0.839$\pm$0.118 | 0.606$\pm$0.175 |
| CLEAR-HPV (raw $h$) | 0.717$\pm$0.138 | 0.832$\pm$0.090 | 0.718$\pm$0.237 | 0.558$\pm$0.233 | 0.828$\pm$0.146 | 0.577$\pm$0.203 |

Table S9: **Logistic regression classifier-based survival prediction on TCGA-HNSCC using CLEAR-HPV.** Results are reported as accuracy (ACC), area under the ROC curve (AUC), F1, precision (Prec), recall (Rec, same as sensitivity), and specificity (Spec) as mean ± 95% confidence interval (CI) across cross-validation folds.
